# Omicron breakthrough infections in vaccinated or previously infected hamsters

**DOI:** 10.1101/2022.05.20.492779

**Authors:** Jie Zhou, Ksenia Sukhova, Paul F. McKay, Ashwini Kurshan, Yeuk Yau, Thomas Lechmere, Jonathan C. Brown, Maya Moshe, Ruthiran Kugasathan, Luke B. Snell, Jonathan D. Edgeworth, Robin J. Shattock, Katie J. Doores, Thomas P. Peacock, Wendy S. Barclay

## Abstract

The second and third years of the SARS-CoV-2 pandemic have been marked by the repeated emergence and replacement of ‘variants’ with genetic and phenotypic distance from the ancestral strains, the most recent examples being Delta and Omicron. Here we describe a hamster contact exposure challenge model to assess protection conferred by vaccination or prior infection against re-infection. We found that 2-doses of self-amplifying RNA vaccine based on the ancestral spike ameliorated weight loss following Delta infection and decreased viral loads, but had minimal effect on Omicron/BA.1 infection. Prior infection with ancestral or Alpha variant was partially protective against Omicron/BA.1 infection, whereas all animals previously infected with Delta and exposed to Omicron became infected, although shed less virus. We further tested whether prior infection with Omicron/BA.1 protected from re-infection with Delta or Omicron/BA.2. Omicron/BA.1 was protective against Omicron/BA.2, but not Delta reinfection, again showing Delta and Omicron have a very large antigenic distance. Indeed, cross-neutralisation assays with human antisera from otherwise immunonaïve individuals (unvaccinated and no known prior infection), confirmed a large antigenic distance between Delta and Omicron. Prior vaccination followed by Omicron or Delta breakthrough infection led to a higher degree of cross-reactivity to all tested variants. To conclude, cohorts whose only immune experience of COVID is Omicron/BA.1 infection may be particularly vulnerable to future circulation of Delta or Delta-like derivatives. In contrast, repeated exposure to antigenically distinct spikes, via infection and or vaccination drives a more cross-reactive immune response, both in hamsters and people.

**One Sentence Summary:** Infection with the Delta and Omicron SARS-CoV-2 variants do not provide cross-protective immunity against reinfection with one another in hamsters.

## INTRODUCTION

Omicron is the most recent SARS-CoV-2 Variant of Concern. Following its timely description by multiple laboratories in Africa (*1*), it spread rapidly displacing the previously circulating Delta variant around the world. At least 5 distinct lineages of Omicron have been described including BA.1 and BA.2 which circulated widely early in 2022, followed in recent months by BA.4 and BA.5 (*2*). All Omicron variants carry an unprecedented number of mutations in their genome including over 30 coding changes in the Spike gene alone. This results in a considerable antigenic distance between Omicron Spike and that of other previous variants, especially Delta (*3*). Thus, it is not unexpected that antibodies induced after vaccination with COVID vaccines, that are based on the Spike of the early Wuhan-hu-1 strain, poorly neutralize the Omicron variant (*4-13*). The lack of cross-neutralization between Omicron and earlier variants likely accounts for the observed high transmission of Omicron in populations that are heavily vaccinated and/or have a high rate of previous infection (*14, 15*).

Here we use a hamster model of SARS-CoV-2 transmission to illustrate Omicron vaccine breakthrough and also a high rate of reinfection by Omicron in animals previously infected with Delta variant. The levels of Omicron virus shedding from vaccinated or previously Delta-infected were compatible with the potential for onwards airborne transmission. We also find that hamsters previously infected with Omicron/BA.1 (BA.1) were not protected against infection with Delta virus but did not become virus positive after exposure to an Omicron/BA.2 (BA.2) isolate. Our *in vivo* findings are paralleled by neutralisation assays with human sera collected from individuals who have recovered following infection with earlier SARS-CoV-2 variants that demonstrate a lack of cross-neutralization between Omicron and Delta. These studies offer an important contribution to risk assessing the potential for variants to escape vaccine control, and to reinfect previously infected individuals. Taken together the results reinforce that there may be specific cohorts who are especially vulnerable to antigenically distant new variants, for example children who have been less vaccinated than adults. Moreover, our findings imply that if we aspire to use vaccines to control circulation of SARS-CoV-2 variants, we will need a system for rapidly updating the immunogen based on detailed antigenic characterization validated with preclinical models.

## RESULTS

### Omicron/BA.1 variant is efficiently transmitted to hamsters vaccinated with a Wuhan-hu-1 Spike saRNA vaccine

We previously showed that immunization with a self-amplifying RNA vaccine encoding the Wuhan-hu-1 SARS-CoV-2 Spike gene (saRNA-Spike) protected hamsters against weight loss following infection through exposure to cage mates infected with either a first wave isolate or an Alpha variant isolate (*16, 17*). Although all exposed immunized hamsters in that study still became virus positive, weight loss and virus shedding were significantly reduced compared to that in a control group vaccinated with an irrelevant immunogen (*17*).

Here we used the same protocol to test the efficacy of the saRNA-Spike vaccine against Delta and BA.1 variants. Groups of 16 hamsters were immunized with either saRNA-Spike or a control vaccine encoding HIV gp120 (saRNA-HIV). The immunization regimen consisted of an initial priming dose, followed by a boosting dose 4 weeks later (Fig. 1A). Two weeks post-boost, serum samples were collected before challenge by the direct contact exposure route. Pseudovirus neutralization assays confirmed that animals vaccinated with saRNA-Spike had high serum neutralizing titres against WT/D614G (ND_50_ = 678, geometric mean), and showed 2-fold decrease (*p* > 0.05) and 13-fold (*p* = 0.0002) decrease in neutralizing activity against Delta and BA.1 variants respectively (Fig. 1B).

**Fig. 1.**
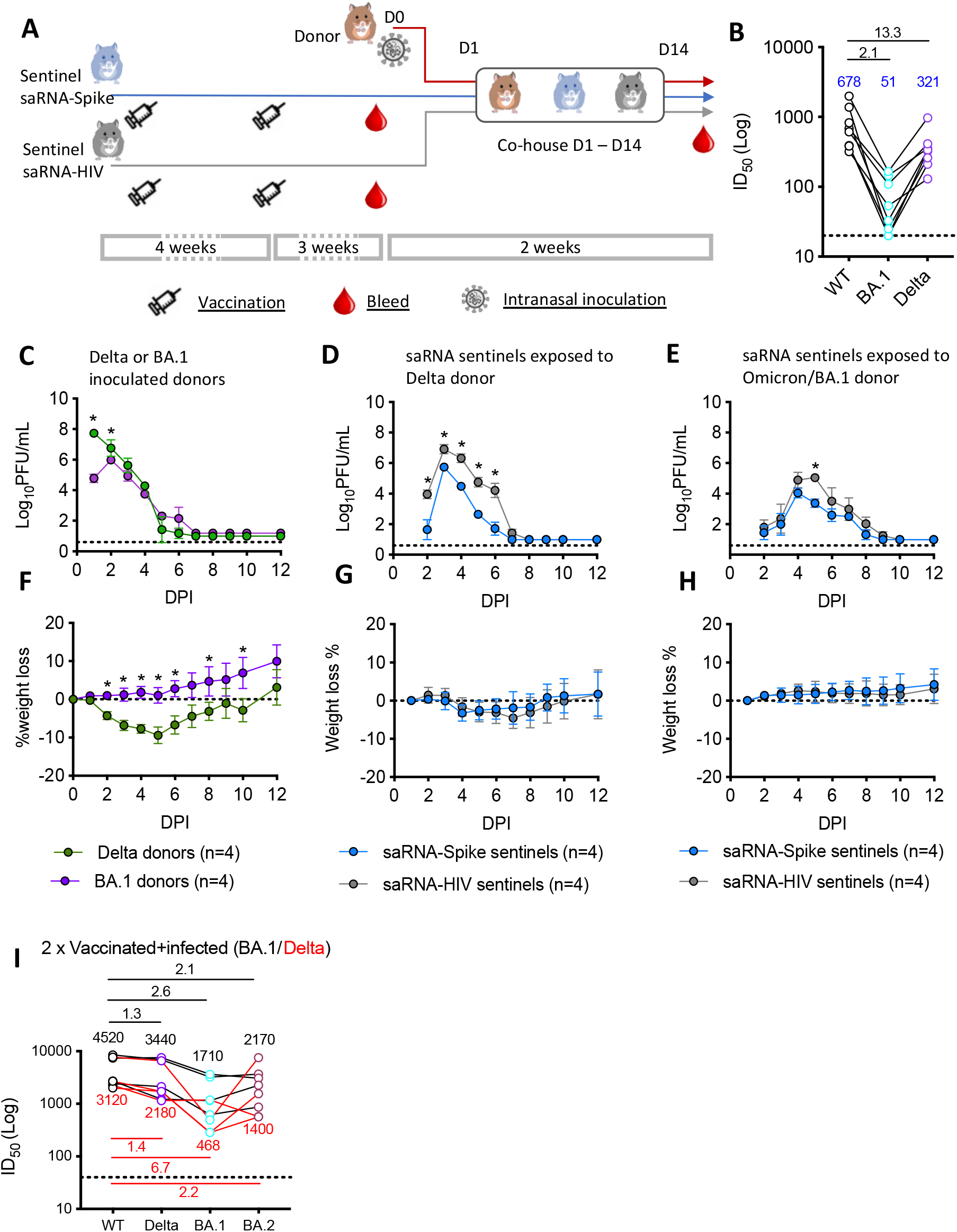
Omicron/BA.1 variant infection of hamsters vaccinated with a Wuhan-like Spike self-amplified RNA vaccine. (**A**) Experimental design. Two groups of 8 hamsters each were vaccinated with self-amplified RNA (saRNA) Spike (n=8) or the saRNA-HIV vaccine (n=8). The vaccination schedule was a priming dose followed 4 weeks later by a boost. Three weeks after the second dose of vaccine, the vaccinated hamsters were co-housed with donor hamsters which had been inoculated intranasally with 100 PFU Delta or BA.1 variant the previous day. Each cage thus housed one donor, one saRNA-Spike vaccinee and one control saRNA-HIV vaccinee hamster. (**B**) Pseudovirus neutralisation assays using sentinel hamster sera collected two weeks after the second vaccine dose. Fold changes (black numbers) in geometric means (blue numbers) are shown above the symbols. (**C-E**), Infectious virus shed in nasal wash of donor hamsters (C), of saRNA sentinels exposed to Delta donors (D), and of saRNA sentinels exposed to the BA.1 donors (E). Nasal wash samples were collected daily and infectious virus titers assessed by plaque assay in Vero cells expressing ACE2 and TMPRSS2 (VAT). Lowest detection limit is 10 PFU/mL. The symbols represent mean and S.D. (**F-H**), Body weight change was monitored daily. Symbols represent mean and S.D. (**I**) Pseudovirus neutralisation assays using sentinel hamster sera collected at the end of experiment, 14 days after exposure. Fold changes in geometric means from the vaccinated sentinels infected with BA.1 (black numbers) or Delta (red numbers) are shown. Statistically significant differences (**C-H**) were determined using Mann-Whitney test. * *p*<0.05, *** *p*<0.001.

Donor hamsters were all productively infected intranasally with 100 PFU of either Delta or BA.1 variant and shed infectious virus in their nasal wash from day 1 post-inoculation. Infectious viral loads in nasal wash were higher on day 1 and day 2 post infection in Delta infected donors than BA.1 infected animals and viral RNA levels were also higher in the nasal wash of Delta infected donors (Fig. 1C and fig. S1A). The Delta inoculated donor hamsters lost approximately 10% starting body weight by day 5 post-infection after which they recovered, while the BA.1 inoculated donor hamsters did not lose weight (Fig. 1F).

Vaccinated hamsters were co-housed with an infected donor 1 day after inoculation. Each cage housed one donor, one saRNA-spike vaccinee and one control saRNA-HIV vaccinee animal (Fig. 1A). Analysis of infectious virus and E gene in nasal washes collected daily revealed that all sentinel hamsters became infected (Fig. 1D, E and fig. S1B, C). However, the infectious viral load and viral RNA copies shed in nasal wash of saRNA-Spike vaccinated hamsters infected with Delta variant was significantly lower than in the control group on every day that virus was detected and in total (area under the curve) (Fig. 1D and fig. S2B). In contrast, infectious viral load in nasal washes of animals infected with Omicron virus was minimally affected by vaccination and was only lower than in control animals on day 5 (Fig 1E and fig. S2C). Sentinels infected by Delta variant lost less weight than the directly infected donor animals regardless of vaccine status; weight loss peaked at around 4.5% in the control group and at 3.2% in the saRNA-Spike vaccine recipients vs 10% in the naïve donors. This likely results from a lower dose initiating infection through the contact exposure route than for the directly inoculated animals. None of the sentinel animals exposed to BA.1 variant in any vaccinated group lost weight (Fig. 1G, H).

We performed pseudovirus neutralization assays with sera collected on day 14 post-infection of vaccinated animals. Neutralizing titres against WT, Delta, BA.1 and BA.2 all increased following breakthrough infection (Fig. 1I). The post-vaccine breakthrough infection sera from the saRNA-Spike vaccinated hamsters infected by BA.1 showed slightly lower neutralizing titres (2.1 and 2.6 fold) against BA.1 and BA.2 than the WT and Delta variants. The post-infection sera from the saRNA-Spike vaccinated hamsters infected with Delta had the lowest neutralizing titres against BA.1, compared to against WT, Delta and BA.2, (6.7-fold lower than against WT) suggesting that vaccine breakthrough with Delta might lead to a greater cross-reactive neutralising response against BA.2, than BA.1.

Thus, the reduction in neutralizing activity induced against BA.1 led to a failure of the vaccine based on first wave Spike sequence to reduce both infection and viral load.

### Reinfection of hamsters infected with earlier variants following exposure to Omicron

We next tested whether BA.1 would also escape prior immunity conferred in animals that were previously infected with earlier variants (Fig. 2A). Groups of 4 hamsters were infected via intranasal inoculation with 100 PFU of an early first wave wildtype isolate (WT/D614G), an Alpha variant isolate, a Delta variant isolate, or were mock infected. All infected animals robustly shed virus in the nasal washes (fig. S3B-G). Six weeks after being infected, serum samples were collected from the recovered animals to test for the presence of neutralizing antibodies. We detected robust neutralizing titres against the homologous Spike within all groups although due to limited serum volumes available we were not able to establish end point titres for all animals. Titres against BA.1 variant fell below the limit of detected in all these earlier variant sera (Fig. 2B-D). The previously infected or naïve hamsters were then co-housed with donor hamsters infected by intranasal inoculation with 100 PFU of BA.1, from day 1 post-infection of the donor. All animals were nasal washed daily. All 4 naïve sentinel animals acquired BA.1 infection from the co-housed donors and shed high levels of BA.1 in their nasal wash (Fig. 2E and fig. S3A). In contrast, only 1 of 4 animals previously infected with first wave virus (WT/D614G) or Alpha variant shed infectious virus in nasal wash after exposure to BA.1-infected donors (Fig. 2F, G and fig. S3H, I). Conversely, no protection against BA.1 infection was observed following prior infection with Delta variant; all 4 animals in that group shed high viral RNA loads and high titres of detectible live virus for several days post-exposure (Fig. 2H and fig. S3J). None of the animals lost weight following the reinfection with BA.1 variant regardless of previous SARS-CoV-2 infections, including the previously mock infected ones (fig. S4).

**Fig. 2.**
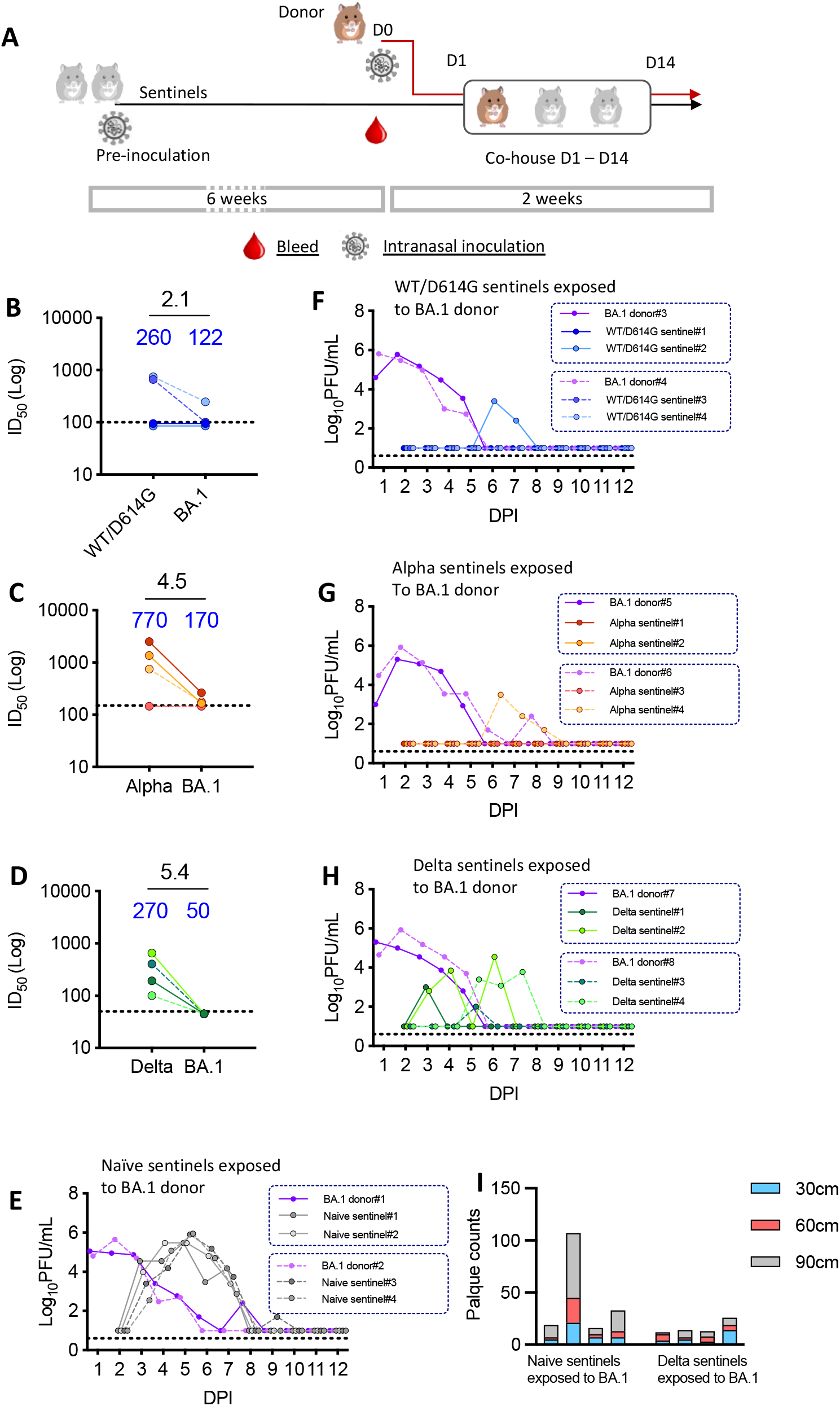
Reinfection of hamsters infected with earlier variants following exposure to Omicron/BA.1. (**A**) Experimental design. Three groups of 4 hamsters each were inoculated intranasally with 100 PFU of WT/D614G, Alpha or Delta variants. Six weeks later, two previously infected hamsters were co-housed with a donor hamster inoculated with 100 PFU BA.1 from 1 day post inoculation. Age-matched naïve sentinels were exposed to the BA.1 donors as controls. Each cage thus had one donor and two sentinels. (**B-D**) Pseudovirus neutralisation assays against homologous strain and BA.1 using sera collected from previously WT/D614G infected hamsters (**B**), Alpha infected hamsters (**C**) and Delta infected hamsters (**D**). Fold changes (black numbers) in geometric means (blue numbers) are shown above the symbols. (**E-H**), Virus shedding profiles of BA.1 infected donors and naïve sentinel hamsters (E), BA.1 infected donors and sentinel hamsters previously infected with WT/D614G (F), BA.1 infected donors and sentinel hamsters previously infected with Alpha (G), and BA.1 infected donors and sentinel hamsters previously infected with Delta (H). Nasal wash samples were collected daily and assessed by the plaque assay. The lower detection limit is 10 PFU/mL. (**I**) Potential for onwards transmission of BA.1 determined by measuring infectious virus deposited from air at 30cm, 60cm and 90cm from the infected naïve and previously Delta infected sentinels.

We also tested the potential of the reinfected animals for onwards transmission by measuring infectious virus exhaled through airborne droplets emitted by the infected animals (fig. S5). At day 2 following the exposure to the infected donor animals, the 4 naïve sentinels and the 4 previous Delta infected hamsters were placed in a chamber from which air was passed over the surface of susceptible cultured cells. Following 10 minutes of exposure, the cells were removed and an overlay applied before 3 day incubation in order to observe plaques formed by infectious virus deposited on the cells. Cell culture plates were placed at 3 different distances, 30 cm, 60 cm or 90 cm from the infected animal. All 8 infected hamsters emitted virus into the air,, and overall plaque counts were not significantly different when comparing those from the naïve sentinels and ones from the animals previously infected with Delta variant (*p* = 0.11) (Fig. 2I). The detection of infectious virus in the nasal washes and in the air emitted from the previously infected animals suggest their potential to support onward chain of transmission despite having prior immunity against SARS-CoV-2.

### Reinfection of hamsters previously infected by Omicron/BA.1 with Delta variant, but not Omicron/BA.2

Due to the low protection conferred by prior infection with Delta variant against BA.1 re-infection, and the fact that both Omicron and Delta variants continue to co-circulate in some parts of the world (albeit now at low levels), we next tested the reciprocal relationship by challenging hamsters previously infected with BA.1 by exposure to the Delta variant. As before, the BA.1 infected hamsters were allowed to recover and convalesce for 6 weeks following the initial infection. Donor animals were infected via intranasal inoculation with 100 PFU of either Delta variant or the second Omicron lineage, BA.2. Sera collected from the previously infected animals at 6 weeks post-infection showed high neutralizing titres against BA.1, two sentinels showed relatively high neutralizing titres against BA.2 Spike, the rest had low or non-detectable titres against BA.2 or Delta Spike (Fig. 3A). Donor hamsters infected with Delta variant lost weight as seen previously while donors infected with BA.2 did not lose weight, similar to the BA.1-infected donors described previously (Fig. 3B). The viral load in nasal washes of BA.2 infected hamsters was lower than that of Delta infected donors (Fig. 3C-E and fig. S6A-C). All 4 Delta-exposed animals were reinfected and shed robust viral E gene loads and infectious virus in nasal washes (Fig. 3E and fig. S6C). 3 of the 4 Naïve age-matched sentinel hamsters exposed to the BA.2 donors were infected, and virus shedding became detectable in their nasal wash samples with a one day delay comparing to the kinetics of transmission measured for BA.1 (see Fig. 2E) (Fig. 3C and fig. S6A). However, only one of the 4 animals previously infected with BA.1 weas reinfected by BA.2 (Fig. 3D and fig. S6B), and titres of infectious virus were low and transient, only evident at 3 dpi. None of the reinfected animals showed weight loss (fig. S7). We also monitored the potential of the animals reinfected with Delta after BA.1 to support onwards airborne transmission. Infectious virus was detected from droplets emitted into the air by these hamsters on day 2 post exposure to the donors (Fig. 3F). Following the exposure to BA.2, neither the directly inoculated donors, nor previously naïve sentinels, nor sentinels previously exposed to BA.1 shed detectible infectious virus particles into the air. Overall, this shows that prior BA.1 infection is partially protective against reinfection with BA.2, but not Delta variant.

**Fig. 3.**
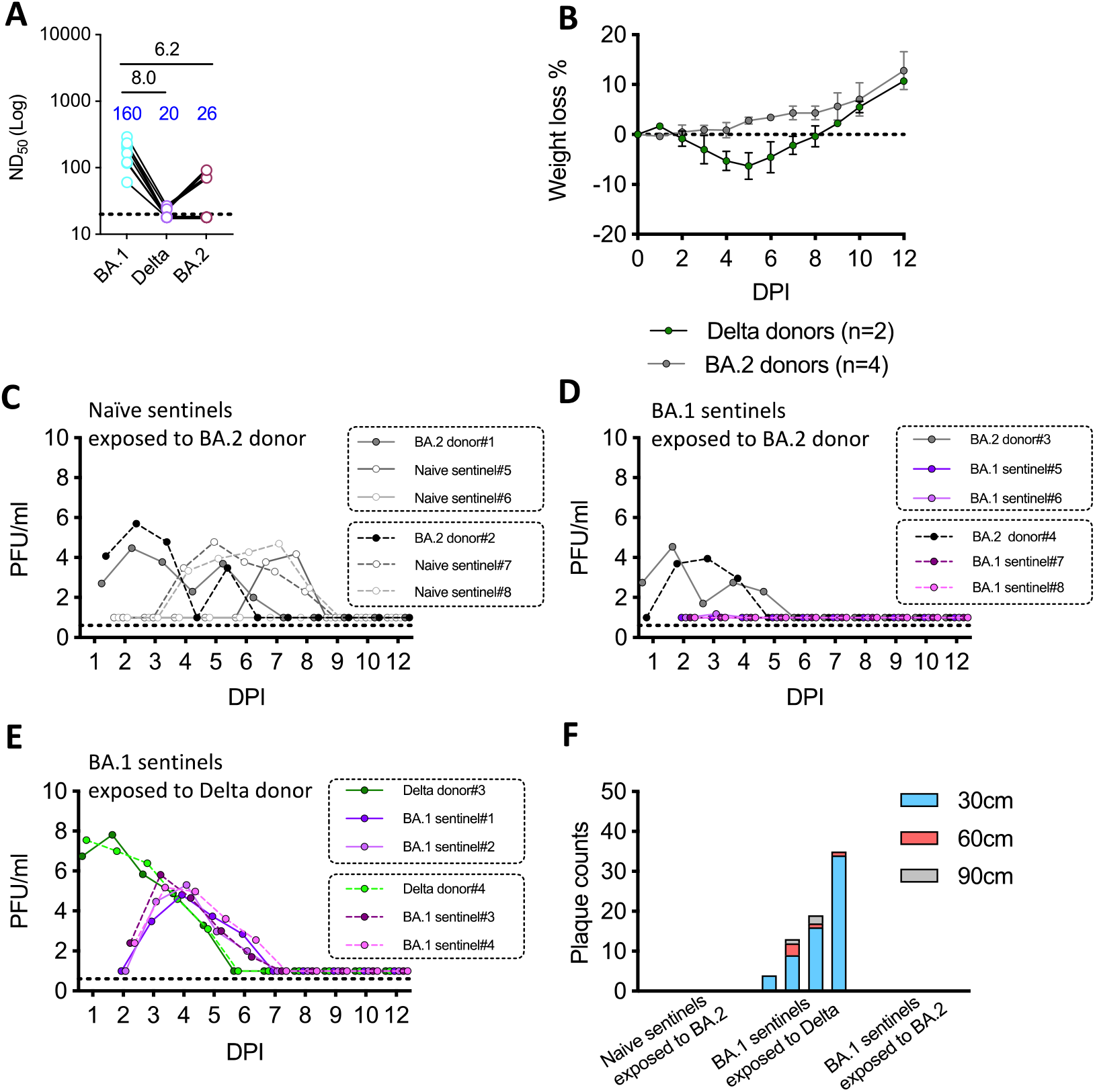
Reinfection of hamsters previously infected by Omicron/BA.1 with Delta variant but not Omicron/BA.2. Eight hamsters were inoculated intranasally with 100 PFU of the BA.1 variant. Six weeks later two previously infected hamsters were co-housed with a donor hamster inoculated with 100 PFU of the Delta or BA.2 variant from 1 day post inoculation. Age-matched naïve sentinels were exposed to BA.2 donors as controls. (**A**) virus shedding in nasal wash samples of the naïve sentinels and the BA.2 donors. (**B**) Potential for onwards transmission by measuring infectious virus deposited at 30cm, 60cm and 90cm from the naïve hamsters infected with Omicron/BA.2 and the pre-BA.1 hamsters reinfected with BA.2 or Delta. (**C**) Pseudovirus neutralisation assays against homologous strain BA.1, Delta and BA.2 using the sera collected from pre-BA.1 infected hamsters. Fold changes (black numbers) in geometric means (blue numbers) are shown above the symbols. (**D)** Virus shedding profile of the Delta donors and pre-BA.1 inoculated sentinels. (**E**) virus shedding profile of the BA.2 donors and pre-BA.1 inoculated sentinels. Nasal wash samples were collected daily and assessed by plaque assay. The lower detection limit of plaque assay is 10 PFU/mL.

### Human convalescent sera

To relate the result obtained from the *in vivo* challenge experiments to the measurable antibody responses in humans, we collected and tested the neutralisation activity of antisera from hospitalised individuals taken during the first UK SARS-CoV-2 wave (N=9), the Alpha wave (N=9), the Delta wave (N=12) and the BA.1 wave (N=16) (Fig. 4). Where possible we chose sera from previously naïve, unvaccinated individuals whose only known exposure was to the named variant. Sera was collected 13-24 days post onset of symptoms. All infected individuals produced a robust neutralizing response against the strain they were initially infected with. Cross-neutralization of other variants was observed but always at reduced titres compared to homologous virus. In sera from first wave, Alpha or Delta infected individuals, the greatest reduction was observed against BA.1. The loss of titre against BA.1 following Delta infection was profound, with a 120-fold reduction in the GMT, compared to the 60-fold reduction in sera from Alpha infected individuals. Sera from individuals infected with WT D614G first wave virus showed a 128.5 fold reduction in ability to neutralize BA.1. In sera from BA.1 infected individuals, a 23-fold reduction and a 15-fold reduction in GMT against Delta and BA.2 respectively were determined in comparison to that against BA.1. These data highlight greater antigenic distance between Omicron and previous variants of concern and also a considerable antigenic distance between BA.1 and BA.2 in otherwise naïve individuals.

**Figure 4.**
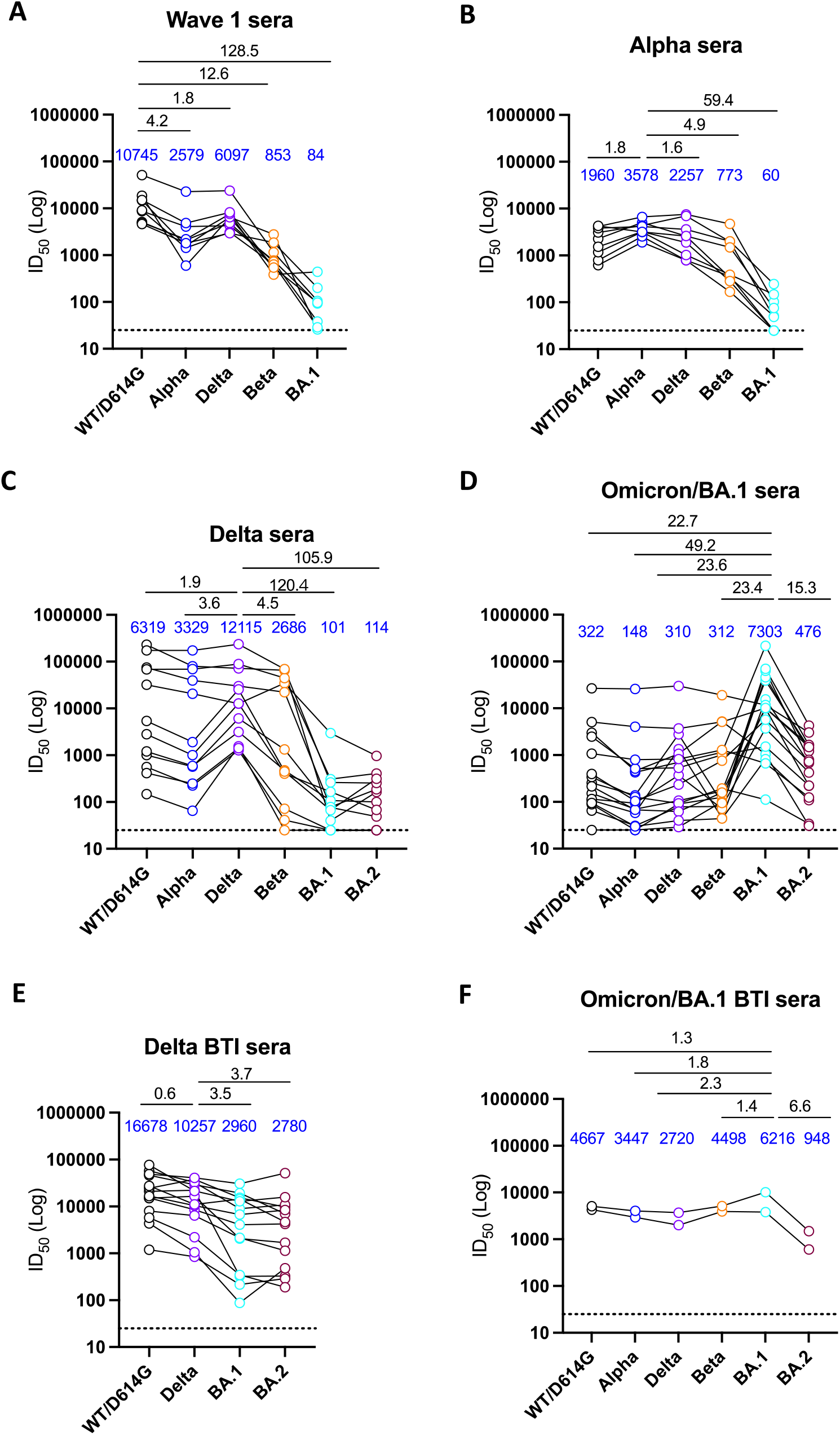
Human convalescent sera. Differences in cross neutralising activity against the variants of concern from the convalescent sera of the individual previously infected with WT/D614G (A), Alpha (B), Delta (C) or BA.1 (D), or vaccinated individuals infected with Delta (E) or BA.1 (F). ID_50_ was measured using HIV-1-based virus particles (PVs), pseudotyped with the S glycoprotein of SARS-CoV-2. Each line represents one individual. The cut-off (dot lines) for the pseudovirus neutralisation assay is 1:50. Fold changes (black numbers) in geometric means (blue numbers) are shown above.

We also measured neutralizing antibody titres in individuals who experienced SARS-CoV-2 infection following 2 prior vaccine doses. These vaccine breakthrough infections yielded antibody titres higher than after vaccination alone, and appeared broader. The drop in antibody titre against BA.1 following Delta infection in vaccinated individuals was only 3.5-fold compared to homologous titres. Interestingly, following vaccine breakthrough infection with BA.1, the largest fold reduction from homologous titres for the two samples was against BA.2, of 6.6-fold.

## DISCUSSION

As SARS-CoV-2 continues to spread at high rates in the human population, the virus is constantly evolving, with new variants periodically emerging. When an older variant is replaced by a new variant, it is vital to understand 1) whether the new variant has a fitness advantage over the old variant; 2) whether the new variant is more transmissible; 3) whether the vaccine’s protection against the new variant decreases 4) whether immunity acquired from prior infections decreases against the new variant. Answering these questions helps us further understand the risk of new variants spreading in a population with ever increasing immunity to SARS-CoV-2. In this study, we used the hamster model to demonstrate that saRNA vaccine encoding Wuhan-hu-1 Spike protein confers significantly reduced neutralising ability against the BA.1 variant (Table 1). Moreover, hamsters previously infected with pre-Omicron variants can be re-infected with Omicron (Table 1) which can be exhaled as infectious virus into the air. These results explain, at least in part, the extremely rapid replacement of the Delta variant by Omicron.

**Table 1.** Omicron breakthrough infections in previously vaccinated or infected hamsters.

| saRNA Vaccine | Re-challenge virus | Infection confirmed<br>by plaque assay | Area under<br>the Curve | Note |
| --- | --- | --- | --- | --- |
| saRNA-Spike | Delta | 4/4 (100%) | 15.5 | Median of (16.5, 12.4, 15.1, 15.9) |
| saRNA-HIV | Delta | 4/4 (100%) | 27.8 | Median of (28.6, 28.5, 24.1, 27.2) |
| saRNA-Spike | Omicron/BA.1 | 4/4 (100%) | 14.3 | Median of (19.8, 14.9, 12.9, 13.7) |
| saRNA-HIV | Omicron/BA.1 | 4/4 (100%) | 21.8 | Median of (27.0, 27.1, 16.7, 14.7) |
| **Primary-infected** |  |  |  |  |
| N/A | Omicron/BA.1 | 4/4 (100%) | 22.7 | Median of (22.2, 23.1, 24.1, 21.2) |
| WT/D614G | Omicron/BA.1 | 1/4 (25%) | 5.8 | Single value |
| Alpha | Omicron/BA.1 | 1/4 (25%) | 5.9 | Single value |
| Delta | Omicron/BA.1 | 4/4 (100%) | 6.6 | Median of (3.0, 11.2, 2.0, 10.3) |
| N/A | Omicron/BA.2 | 3/4 (75%) | 17.6 | Median of (8.0, 17.6, 18.8) |
| Omicron/BA.1 | Omicron/BA.2 | 1/4 (25%) | 1.0 | Single value |
| Omicron/BA.1 | Delta | 4/4 (100%) | 16.20 | Median of (14.9, 14.8, 17.6, 18.7) |

In our previous study, we showed that the saRNA-Spike vaccine confers protections against the WT/D614G and Alpha variants after two vaccine doses following the challenge of vaccinated hamsters via a direct contact route. In this study, fully saRNA vaccine-immunised hamsters had decreased virus titres and weight loss after co-housing with Delta inoculated donor hamsters. The saRNA vaccine introduced crossing neutralizing activity against the Delta variant, but the neutralising ability against BA.1 was significantly reduced. We did not observe a difference in virus shedding profiles or weight loss between the saRNA-vaccinated group and the control group, although other hamster experiments had also demonstrated that BA.1 did not cause any weight loss (*18-20*). Doremalen et al. also showed that upon vaccination with AZD1222 (ChAdOx1, a replication-deficient simian adenovirus-vectored vaccine encoding the Spike protein), antibody titres dropped significantly against the Omicron variant, and the AZD1222 vaccinated hamsters inoculated intranasally with BA.1 had similar shedding on day 1 and day 2 compared to controls (*21*). This is consistent with the real-world survey, where breakthrough infections with Omicron are more likely to occur in people who are fully vaccinated, compared to the previous variants. In a Danish household study, fully vaccinated people experienced higher secondary attack rates in households with Omicron compared to Delta (*14*). Many groups have reported that the Omicron spike evades neuralization by antisera from convalescent patients or individuals vaccinated with two doses of mRNA vaccine (*4-13*). These data suggest that immune evasiveness is likely largely responsible for the rapid spread of the Omicron variant. We also showed that both Delta and Omicron breakthrough infection in double saRNA-Spike vaccinated hamsters generated potent and broad neutralizing activity against the current variants of concern, including Omicron BA.1 and BA.2, which is consistent with the data presented here and the report by Lechmere et al (*22*), suggesting that breakthrough SARS-CoV-2 infection can generate widely cross-reactive antibodies.

More and more people have experienced COVID-19 more than once since the beginning of the pandemic. Infection can induce strong protection against reinfection with Alpha, Beta and Delta variants (*23-26*). However, the effectiveness of previous infection in preventing reinfection was estimated to be 56.0% against the Omicron (*27*). Our findings suggest that previous infections with the WT/D614G, Alpha and Delta variants can confer cross-protection against the Omicron by reducing infection, virus peak titres, and virus shedding duration. Although our results are based on a small number of animals, the protection gained from previous Delta infection was worse against BA.1, when compared to the protection from previous infections with WT/D614G or Alpha. This is consistent with our and other serology studies based on a unique set of sera from convalescent patients infected with a range of variants (*26*). In an antigenic map created by Straten et al., the WT/D614G and Alpha variants group together. Infection by the WT/D614G and Alpha induced broader and stronger immunity compared to Delta. Understanding the antigenicity of SARS-CoV-2 Spike is essential for risk assessment of re-infection as well as strain selection for COVID-19 vaccine updates. In addition, we confirmed that previously Delta-infected sentinels reinfected with Omicron can exhale infectious virus into the air, and thus confer onward airborne transmission. Unlike the previous variants, Omicron infection generally causes less severe disease, and many reinfections are asymptomatic. (*26, 28*) Viral shedding from these infected and reinfected individuals that are asymptomatic (or mildly symptomatic) could pose a serious public health concern to the unvaccinated or immunocompromised populations. We conclude that high reinfection rates and potential airborne transmission of reinfected individuals contribute to the rapid spread of the Omicron variant.

At present every new SARS-CoV-2 variant has arisen from a pre-variant ancestor. However, it is hypothesized that future variants could arise from previous variants such as Alpha or Delta. Here we have shown that prior BA.1 infection provides very poor protection against Delta reinfection (and vice versa). Therefore, although Delta cases are now at very low levels globally, Delta could potentially have an advantage in populations which have suffered high burdens of Omicron and have very low vaccine rates, such as young children. Furthermore, a future variant derived from Delta but which has evolved orthogonal antigenicity to both ancestral strains and Omicron, could have a large advantage in populations which have had both high vaccine rates and high levels or prior infections with Omicron.

The BA.2 lineage has been circulating in populations since the start of 2022, and currently BA.2 is estimated to account for over 95% cases in England (*15*). A few cases of sequence-confirmed reinfections with BA.2 following BA.1 have been detected (*29, 30*). Mykytyn et al suggested that Omicron BA.1 and BA.2 have evolved as two distinct antigenic outliers (*31*). We therefore assessed the effectiveness of BA.1 infection against reinfection with BA.2 using the hamster direct contact challenge model. Unlike other variants, BA.2 replicated to lower titres in directly inoculated naïve hamsters, and did not transmit efficiently via direct contact route. Antisera collected from the BA.1 infected hamsters showed a significant reduced neutralising activity against the BA.2, as well as Delta. In contrast to 100% reinfection in the Omicron sentinels exposed to the Delta donors, only one BA.1 sentinel hamster was reinfected with BA.2 and transiently shed virus. Although Omicron BA.1 infection does not introduce a robust antibody response against BA.2, it still prevented BA.2 reinfection via direct contact route in this model.

In conclusion, our study emphasises the utility of the hamster model in studying vaccine efficacy and the potential reinfection with emerging SARS-CoV-2 variants. Our findings provide insights into the rapid surge of the Omicron variant, and this information will be important for making evidence-based public health policies.

## MATERIALS AND METHODS

### Biosafety and ethics statement

All work performed was approved by the local genetic manipulation (GM) safety committee of Imperial College London, St. Mary’s Campus (centre number GM77), and the Health and Safety Executive of the United Kingdom, under reference CBA1.77.20.1. Animal research was carried out under a United Kingdom Home Office License, P48DAD9B4.

Collection of surplus serum samples was approved by South Central – Hampshire B REC (20/SC/0310). SARS-CoV-2 cases were diagnosed by RT-PCR of respiratory samples at St Thomas’ Hospital, London.

### Cells and viruses

Human embryonic kidney cells (293T; ATCC; ATCC CRL-11268) were maintained in Dulbecco’s modified Eagle’s medium (DMEM; Gibco), 10% fetal calf serum (FCS), 1x non-essential amino acids (NEAA; Gibco), 1x penicillin-streptomycin (P/S; Gibco). Stably transduced ACE2-expressing 293T cells were produced as previously described (*32*), and maintained with the addition of 1 μg/ml puromycin to growth medium. African green monkey kidney (VeroE6) cells expressing human angiotensin-converting enzyme 2 (ACE2) and transmembrane protease serine 2 precursor (TMPRSS2) (VeroE6-ACE2-TMPRSS2) were kindly provided by MRC-University of Glasgow Centre for Virus Research (CVR), Glasgow (*33*). The cells were maintained in DMEM, 10% FCS, 1 mg/mL Geneticin (Gibco), 0.2 mg/mL Hygromycin B (Invitrogen). All viral stocks used in this study were grown in the VeroE6-ACE2-TMPRSS2 cells.

| Name in text | Virus name | PANGO lineage | GISAID/COG Accession |
| --- | --- | --- | --- |
| WT/D614G | hCoV-19/England/IC19/2020 | B.1.13 | EPI_ISL_475572 |
| Alpha | hCoV-19/England/205080610/2020 | B.1.1.7 | EPI_ISL_723001 |
| Delta | hCoV-19/England/SHEF-10E8F3B/2021 | B.1.617.2 | EPI_ISL_1731019 |
| BA.1 | M21021166 | BA.1 | N/A |
| BA.2 | IC243335 | BA.2 | N/A |

### Plaque assays

Nasal wash samples were serially diluted in DMEM and added to the VeroE6-ACE2-TMPRSS2 cell monolayers for 1h at 37°C. Inoculum was then removed and cells were overlayed with DEMEM containing 0.2% w/v bovine serum albumin (Gibco), 0.16% w/v NaHCO_3_ (Gibco), 10 mM HEPES (Invitrogen), 2 mM L-Gutamine (Gibco), 1 X P/S and 0.6% w/v Avicel (Gibco). Plates were incubated at 37°C, 5% CO_2_ for 3 days. The overlay was then removed, and monolayers were stained with 0.05% crystal violet solution for 1h at room temperature. Plates were washed with tap water then dried and virus plaques were counted. The lower limit of detection of the assay was 10 plaque forming units per mL.

### SARS-CoV-2 E gene Real-time RT-PCR

Virus genomes were quantified by Envelop (E) gene RT-qPCR as previously described (*34*). Viral RNA was extracted from supernatants of hamster nasal wash samples using the QIAsymphony DSP Virus/Pathogen Mini Kit on the QIAsymphony instrument (Qiagen). Real time RT-qPCR was then performed using the AgPath RT-PCR (Life Technologies) kit on a QuantStudioTM 7 Flex Real-Time PCR System with the primers specific for SARS-CoV-2 E gene (*35*). For absolutely quantification of E gene RNA copies, a standard curve was generated using dilutions viral RNA of known copy number. E gene copies per ml of original virus supernatant were then calculated using this standard curve. The lower limit of detection of the E gene RT-qPCR was 1200 E copies per mL.

### Hamster transmission studies

Hamster transmission studies were performed in a containment level 3 laboratory, using ISO Rat900 Individually Ventilated Cages (IVC) (Techniplast, U.K). Outbred Syrian Hamsters (4-6 weeks old), weighing 80-130 g were used. In the vaccine study, sentinel hamsters were immunized twice, four weeks apart with an saRNA vaccine encoding either SARS-CoV-2 Spike protein or a control vaccine encoding HIV gp120 protein, intramuscularly in 100 μl. Donor hamsters were intranasally inoculated with 50μl of 100 PFU of each virus while lightly anaesthetised with isoflurane. The vaccinated sentinel hamsters were introduced into the same cage as an infected donor day 1 after inoculation. Each cage thus housed one donor, one saRNA SARS-CoV-2 S vaccinee and one control saRNA HIV gp120 vaccinee animal. In the reinfection studies, sentinel hamsters were intranasally inoculated with 100 PFU of virus. Six weeks later, two pre-infected sentinel hamsters were introduced into the same cage as an infected donor day 1 post inoculation. Each cage thus housed one donor and two sentinel hamsters. Co-house continued to the end of experiments. All animals were nasal washed daily by instilling 400 μl of PBS into the nostrils, the expectorate was collected into disposable 50 ml falcon tubes. Hamsters were weighed daily post-infection.

The potential for hamsters infected with SARS-CoV-2 to transmit onwards was assessed using a set of equipment which detects infectious virus exhaled from infected animals as described previously (*36*). Airflow of 4.5 L/minute was introduced using the bias flow pump via three ports into a 10 cm (height) x 9 cm (diameter) hamster chamber (1.5 L/minute into each port). Sentinel cell culture plates were placed at 3 different distances, 30cm, 60cm or 90cm from the infected animal source.

### Pseudovirus neutralization assays (Imperial College London)

SARS-CoV-2 spike-bearing lentiviral pseudotypes (PV) were generated as described previously (*32, 37*). Pseudovirus neutralization assays were performed by incubating serial dilutions of heat-inactivated human convalescent antisera with a set amount of pseudovirus. Antisera/pseudovirus mix was then incubated at 37°C for 1 h then overlayed into 96 well plates of 293T-ACE2 cells. 48 h later cells were lysed with reporter lysis buffer (Promega) and assays were read on a FLUOstar Omega plate reader (BMF Labtech) using the Luciferase Assay System (Promega).

### Neutralisation assay with SARS-CoV-2 pseudotyped virus (King’s College London)

Pseudotyped HIV-1 virus incorporating the SARS-CoV-2 Spike protein (either wild-type, B.1.1.7, B.1.351, B.1.617.2 or B.1.1.529, BA.2) were prepared as previously described (*38, 39*). Viral particles were produced in a 10 cm dish seeded the day prior with 5×106 HEK293T/17 cells in 10 ml of complete Dulbecco’s Modified Eagle’s Medium (DMEM-C, 10% FBS and 1% Pen/Strep) containing 10% (vol/vol) foetal bovine serum (FBS), 100 IU/ml penicillin and 100 μg/ml streptomycin. Cells were transfected using 90 μg of PEI-Max (1 mg/mL, Polysciences) with: 15μg of HIV-luciferase plasmid, 10 μg of HIV 8.91 gag/pol plasmid and 5 μg of SARS-CoV-2 spike protein plasmid.(*40, 41*) The supernatant was harvested 72 hours post-transfection. Pseudotyped virus particles was filtered through a 0.45μm filter, and stored at -80°C until required.

Serial dilutions of serum samples (heat inactivated at 56°C for 30mins) were prepared with DMEM media (25μL) (10% FBS and 1% Pen/Strep) and incubated with pseudotyped virus (25μL) for 1-hour at 37°C in half-area 96-well plates. Next, Hela cells stably expressing the ACE2 receptor were added (10,000 cells/25μL per well) and the plates were left for 72 hours. Infection levels were assessed in lysed cells with the Bright-Glo luciferase kit (Promega), using a Victor™ X3 multilabel reader (Perkin Elmer). Each serum sample was run in duplicate and was measured against the five SARS-CoV-2 variants within the same experiment using the same dilution series.

### Virus sequencing

Delta or BA.1 variant infection were confirmed using whole genome sequencing as previously described (*38*) or using MT-PCR (*42*).

### Statistical analysis

Statistical analysis was performed using Graphpad Prism. Two-group comparisons were tested using Mann-Whitney test for unpaired groups and Wilcoxon matched-pairs signed rank test was used for paired groups. For all tests, a value of *p* < 0.05 was considered significant.

## List of Supplementary Materials

**Fig. S1.**
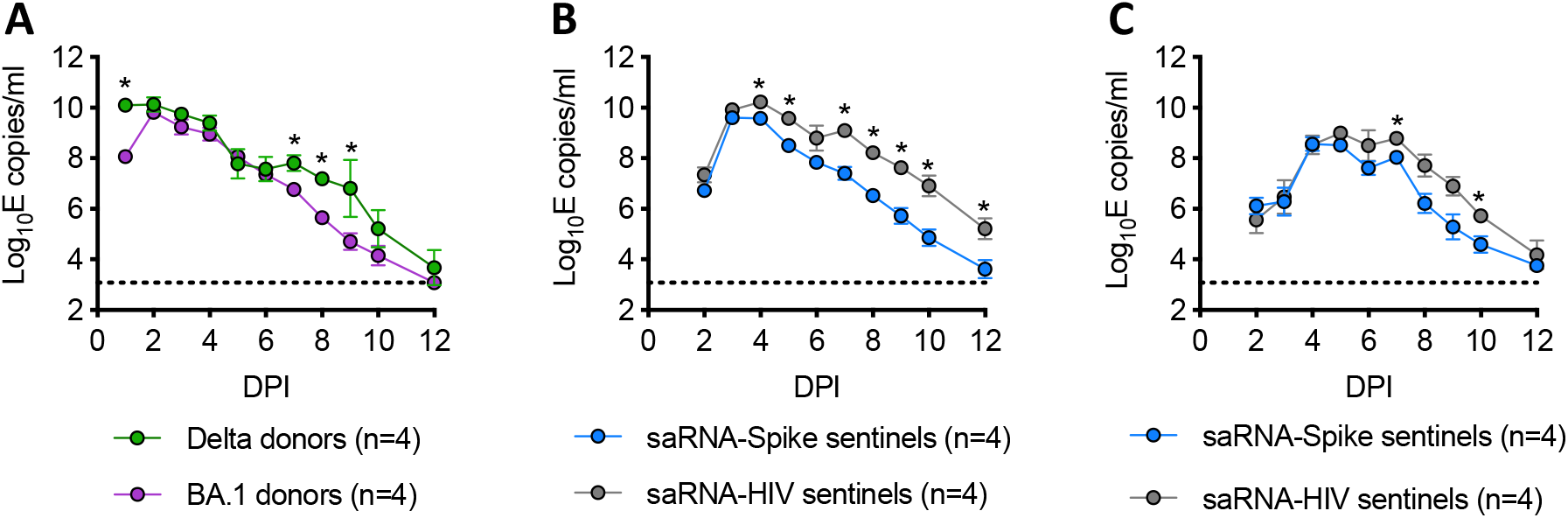
Virus shedding profile of the donor and self-amplifying RNA vaccine (saRNA) sentinel hamsters. **(A)** Virus shedding profile of donor hamsters inoculated with 100 PFU Delta or BA.1. **(B)** Virus shedding profile of saRNA sentinels exposed to the delta donors. (**C)** Virus shedding profile of saRNA sentinels exposed to the BA.1 donors. Nasal wash samples were collected daily and assessed by real-time RT-PCR targeting Envelop gene of SARS-CoV-2. The lower detection limit is 1200 E copies/mL. Statistically significant differences were determined using Mann-Whitney test. * *p*<0.05.

**Fig. S2.**
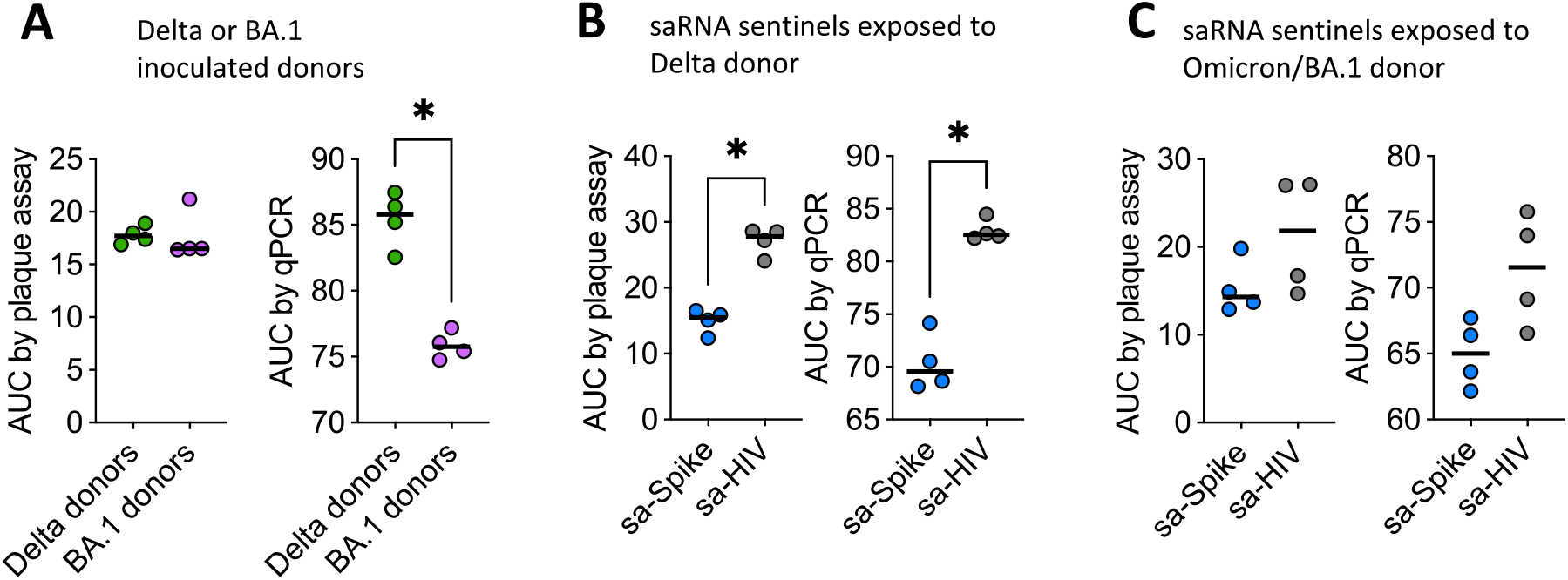
Total virus shedding of the donor and self-amplifying RNA vaccine (saRNA) sentinel hamsters. (**A)** Area under the curve (AUC) of donor hamsters inoculated with 100 PFU Delta or BA.1 was determined by plaque assay or real-time RT-PCR. (**B)** AUC of saRNA sentinels exposed to the delta donors. (**C)** AUC of saRNA sentinels exposed to the Omicron/BA.1 donors. Statistically significant differences were determined using Mann-Whitney test. * *p*<0.05.

**Fig. S3.**
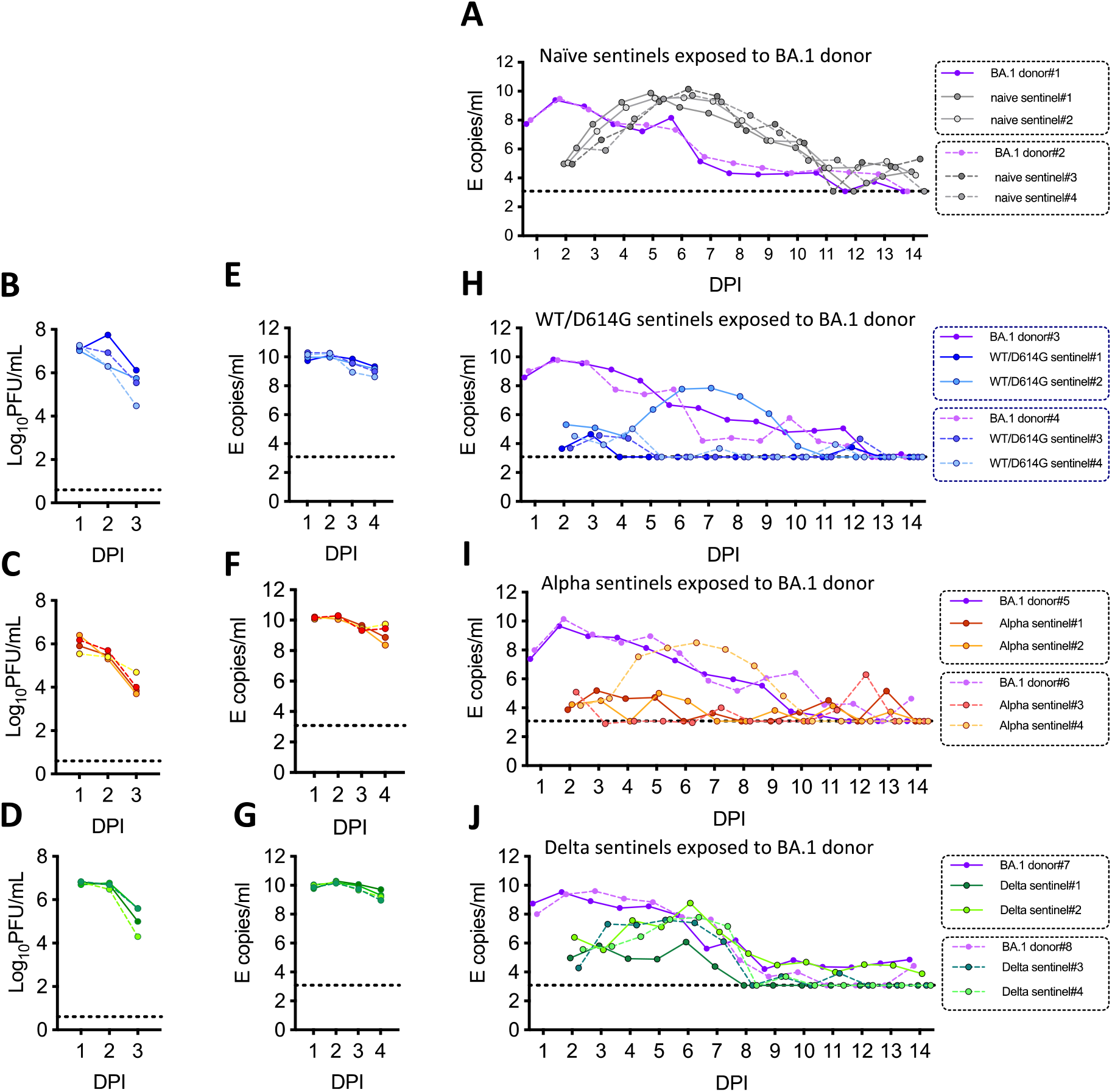
Virus shedding profile of donor and sentinel hamsters following exposure after previous infection. **(A)** Virus shedding profile of BA.1 inoculated donors and naïve sentinel hamsters. **(B-G)**, Hamsters were previously infected with 100 PFU WT/D614G, Alpha or Delta variants. Virus shedding profiles determined by plaque assays **(B–D)** or real-time PCR **(E–G)** from 1 to 3 days post inoculation are shown. (**H-J)** Virus shedding profile of previously infected sentinel hamsters exposed to Omicron/BA.1 donors. Nasal wash samples were collected daily and assessed by real-time RT-PCR targeting E gene of SARS-CoV-2. The lower detection limit is 1200 E gene copies/mL.

**Fig. S4.**
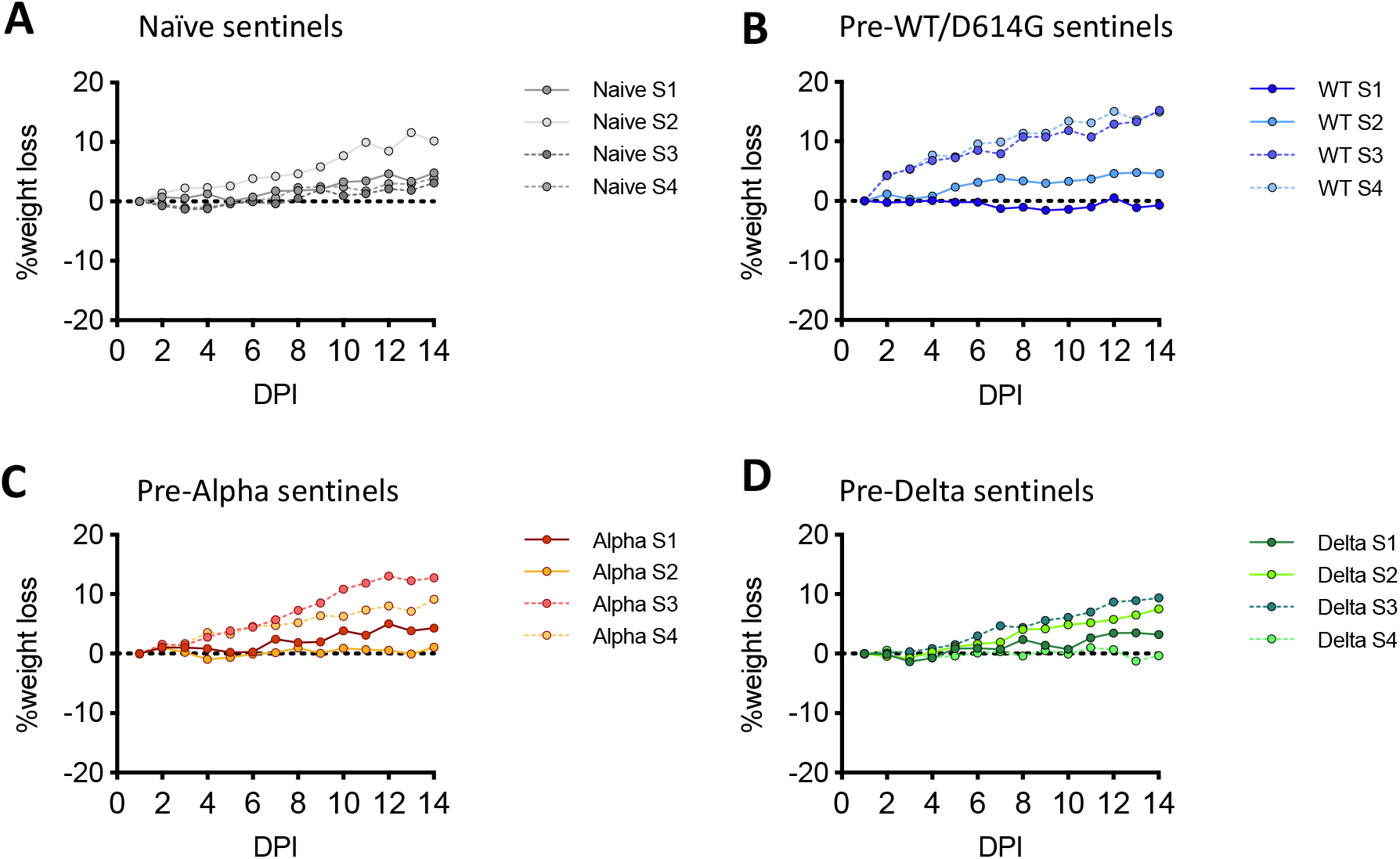
Weight loss change of the sentinel hamsters exposed to Omicron/BA.1 donors. **(A)** Weight loss change of naïve sentinel hamsters exposed to the BA.1 donors. **(B)** Weight loss change of pre-WT/D614G sentinel hamsters exposed to the BA.1 donors. **(C)** Weight loss change of pre-Alpha sentinel hamsters exposed to the BA.1 donors. **(D)** Weight loss change of pre-Delta sentinel hamsters exposed to the BA.1 donors

**Fig. S5.**
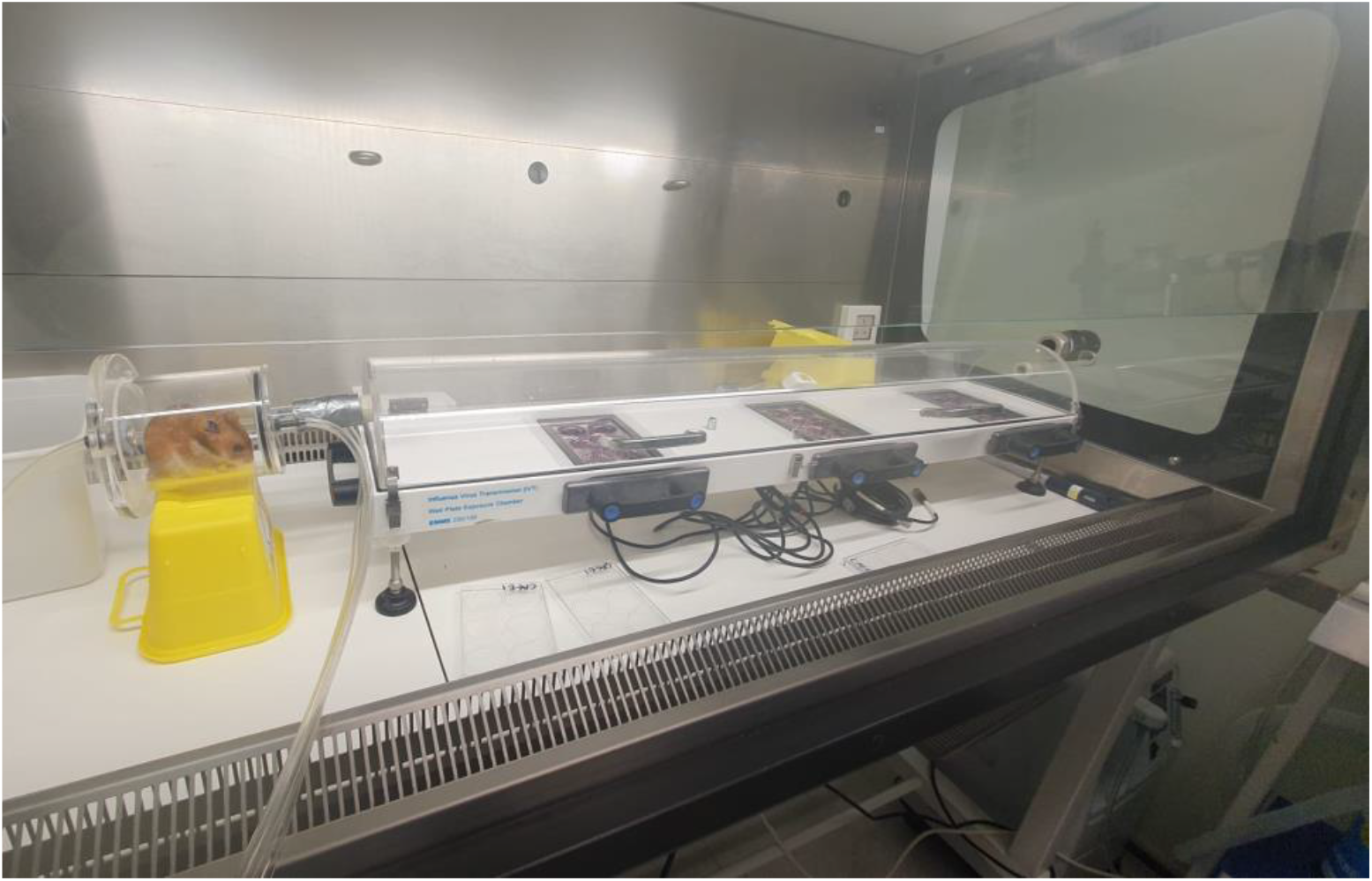
Assessing infectious virus exhaled from infected hamster. Airflow is generated using a bias flow pump, which connects to a hamster chamber (10cm x 9cm, long x diameter). The chamber is connected to a half cylindrical clear acrylic 100cm (length) x 18cm (width) x 9cm (height) exposure tunnel containing cell culture plates situated 30cm, 60cm and 90cm from the animal.

**Fig. S6.**
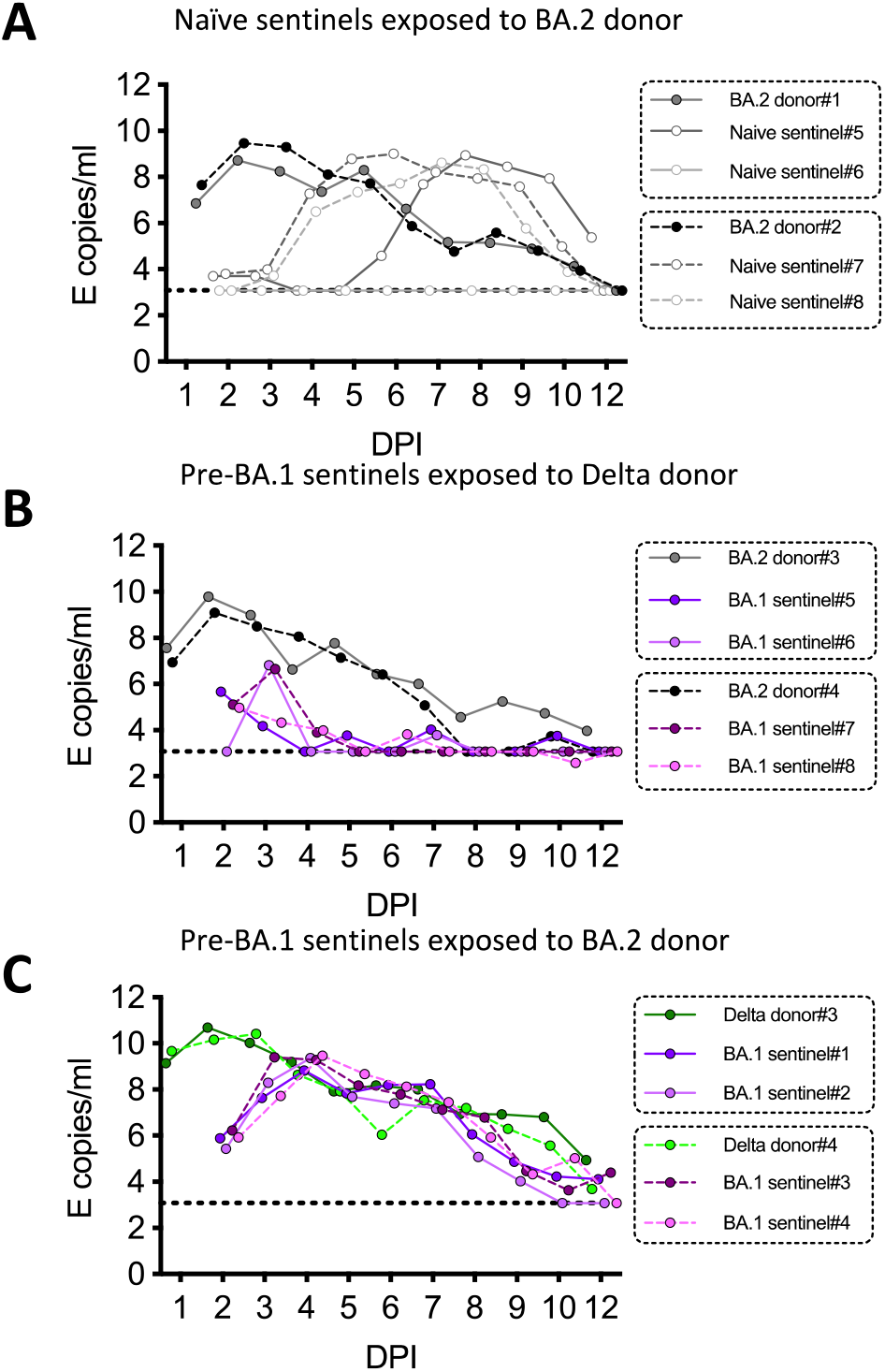
Virus shedding profile of the donor and sentinel hamsters. (**A**) Virus shedding profile of naïve sentinel hamsters exposed to the BA.2 inoculated donors. (**B**) Virus shedding profile of pre-BA.1 sentinel hamsters exposed to BA.2 donors. (**C**) Virus shedding profile of pre-BA.1 sentinel hamsters exposed to Delta donors. Nasal wash samples were collected daily and assessed by real-time RT-PCR targeting Envelop gene of SARS-CoV-2. The lower detection limit is 1200 E copies/mL.

**Fig. S7.**
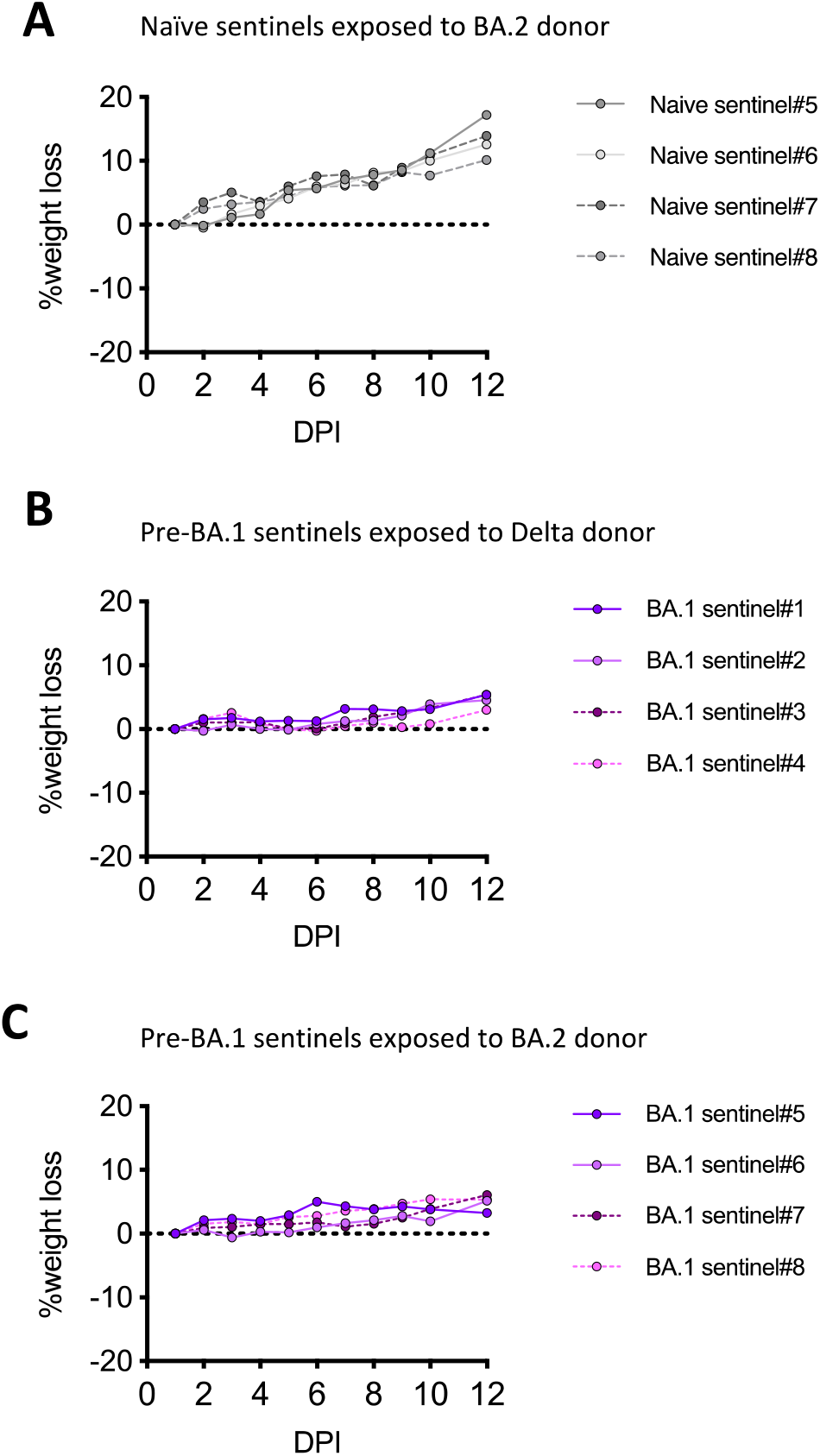
Weight loss change of hamsters infected by Omicron/BA.2 and previously infected sentinel hamsters exposed to Delta or Omicron/BA.2 virus. (**A**) Weight loss of naïve sentinel hamsters exposed to BA.2 donors. (**B**) Weight loss of pre-BA.1 sentinel hamsters exposed to the Delta donors. (**C**) Weight loss change of pre-BA.1 sentinel hamsters exposed to the BA.2 donors.

## Acknowledgements

We thank staff at Imperial College Central Biological Services for their expert help. We thank Dr Matthew Turnbull and Dr Suzannah Rihn of the MRC-University of Glasgow Centre for Virus Research (CVR) for sharing their Vero E6-ACE2-TMPRSS2 cells and Gavin Screaton, Wanwisa Dejnirattisai and Alison Cowper from Oxford University for sharing the BA.1 isolate. For the Alpha, Delta and Omicron swabs, we thank Thushan de Silva at University of Sheffield, the ISARIC4C consortium and Paul Randell, Marcus Pond and colleagues at NWLP and PHE, Michael Crone and Graham Taylor of Imperial College London.

## Funding

This work was supported by the G2P-UK National Virology Consortium funded by the MRC (MR/W005611/1).

RK was supported by Wellcome fellowship no. 216353/Z/19/Z.

## Author contributions

W.S.B., T.P.P., J.Z., K.J.D., and R.J.S. conceptualized the studies; J.Z., K.S., P.F.M., and

M.M. performed the hamster experiments and analyzed the data; T.P.P., A.K., Y.Y., T.L., L.B.S., and K.J.D. performed the neutralization assays and virus sequencing; J.C.B., J.Z., and

M.M. cultured the variants of SARS-CoV-2; W.S.B., J.Z., T.P.P., K.J.D. and K.S. drafted the manuscript, all authors had the opportunity to provide feedback for the final manuscript.

## Competing interests

Robin Shattock and Paul McKay and are co-inventors on a patent application covering this SARS-CoV-2 self-amplifying RNA vaccine. All other authors have nothing to declare.

## Data and materials availability

All data are available in the main text or the supplementary materials.

## References

1. R. Viana et al., Rapid epidemic expansion of the SARS-CoV-2 Omicron variant in southern Africa. Nature 603, 679–686 (2022).

2. H. Tegally et al., Continued Emergence and Evolution of Omicron in South Africa: New BA.4 and BA.5 lineages. medRxiv, 2022.2005.2001.22274406 (2022).

3. A. Netzl et al., Analysis of SARS-CoV-2 Omicron Neutralization Data up to 2021-12-22. bioRxiv, 2021.2012.2031.474032 (2022).

4. N. Andrews et al., Covid-19 Vaccine Effectiveness against the Omicron (B.1.1.529) Variant. New England Journal of Medicine, (2022).

5. S. Collie, J. Champion, H. Moultrie, L.-G. Bekker, G. Gray, Effectiveness of BNT162b2 Vaccine against Omicron Variant in South Africa. New England Journal of Medicine 386, 494–496 (2021).

6. W. Dejnirattisai et al., Reduced neutralisation of SARS-CoV-2 omicron B. 1.1. 529 variant by post-immunisation serum. The Lancet 399, 234–236 (2022).

7. S. Cele et al., Omicron extensively but incompletely escapes Pfizer BNT162b2 neutralization. Nature 602, 654–656 (2022).

8. J. Newman et al., Neutralising antibody activity against SARS-CoV-2 variants, including Omicron, in an elderly cohort vaccinated with BNT162b2. medRxiv, 2021.2012.2023.21268293 (2021).

9. J. M. Carreño et al., Activity of convalescent and vaccine serum against SARS-CoV-2 Omicron. Nature 602, 682–688 (2022).

10. M. Hoffmann et al., The Omicron variant is highly resistant against antibody-mediated neutralization: Implications for control of the COVID-19 pandemic. Cell 185, 447–456.e411 (2022).

11. E. Cameroni et al., Broadly neutralizing antibodies overcome SARS-CoV-2 Omicron antigenic shift. Nature 602, 664–670 (2022).

12. D. Planas et al., Considerable escape of SARS-CoV-2 Omicron to antibody neutralization. Nature 602, 671–675 (2022).

13. Y. Cao et al., Omicron escapes the majority of existing SARS-CoV-2 neutralizing antibodies. Nature 602, 657–663 (2022).

14. F. P. Lyngse et al., SARS-CoV-2 Omicron VOC Transmission in Danish Households. medRxiv, 2021.2012.2027.21268278 (2021).

15. U. H. S. Agency, “SARS-CoV-2 variants of concern and variants under investigation in England.”

16. P. F. McKay et al., Self-amplifying RNA SARS-CoV-2 lipid nanoparticle vaccine candidate induces high neutralizing antibody titers in mice. Nature Communications 11, 3523 (2020).

17. R. Frise et al., A Self-Amplifying RNA Vaccine Protects Against SARS-CoV-2 (D614G) and Alpha Variant of Concern (B. 1.1. 7) in a Transmission-Challenge Hamster Model.

18. R. Suzuki et al., Attenuated fusogenicity and pathogenicity of SARS-CoV-2 Omicron variant. Nature 603, 700–705 (2022).

19. P. J. Halfmann et al., SARS-CoV-2 Omicron virus causes attenuated disease in mice and hamsters. Nature 603, 687–692 (2022).

20. T. P. Peacock et al., The altered entry pathway and antigenic distance of the SARS-CoV-2 Omicron variant map to separate domains of spike protein. bioRxiv, 2021.2012.2031.474653 (2022).

21. D. Neeltje van et al., Efficacy of ChAdOx1 vaccines against SARS-CoV-2 Variants of Concern Beta, Delta and Omicron in the Syrian hamster model. Nature Portfolio, (2022).

22. T. Lechmere et al., Broad Neutralization of SARS-CoV-2 Variants, Including Omicron, following Breakthrough Infection with Delta in COVID-19-Vaccinated Individuals. mBio 0, e03798–03721.

23. H. Chemaitelly, R. Bertollini, L. J. Abu-Raddad, Efficacy of Natural Immunity against SARS-CoV-2 Reinfection with the Beta Variant. N Engl J Med 385, 2585–2586 (2021).

24. L. J. Abu-Raddad et al., Introduction and expansion of the SARS-CoV-2 B.1.1.7 variant and reinfections in Qatar: A nationally representative cohort study. PLoS Med 18, e1003879 (2021).

25. P. Kim, S. M. Gordon, M. M. Sheehan, M. B. Rothberg, Duration of SARS-CoV-2 Natural Immunity and Protection against the Delta Variant: A Retrospective Cohort Study. Clin Infect Dis, (2021).

26. N. Wolter et al., Early assessment of the clinical severity of the SARS-CoV-2 omicron variant in South Africa: a data linkage study. Lancet 399, 437–446 (2022).

27. H. N. Altarawneh et al., Protection against the Omicron Variant from Previous SARS-CoV-2 Infection. N Engl J Med, (2022).

28. P. England, SARS-CoV-2 variants of concern and variants under investigation in England. Tech Brief 12, (2021).

29. U. H. S. Agency, “Risk assessment for SARS-CoV-2 variant: VUI-22JAN-01 (BA.2),” (2022).

30. M. Stegger et al., Occurrence and significance of Omicron BA.1 infection followed by BA.2 reinfection. medRxiv, 2022.2002.2019.22271112 (2022).

31. A. Z. Mykytyn et al., Omicron BA.1 and BA.2 are antigenically distinct SARS-CoV-2 variants. bioRxiv, 2022.2002.2023.481644 (2022).

32. T. P. Peacock et al., The furin cleavage site in the SARS-CoV-2 spike protein is required for transmission in ferrets. Nat Microbiol 6, 899–909 (2021).

33. S. J. Rihn et al., A plasmid DNA-launched SARS-CoV-2 reverse genetics system and coronavirus toolkit for COVID-19 research. PLoS Biol 19, e3001091 (2021).

34. J. Zhou et al., Investigating Severe Acute Respiratory Syndrome Coronavirus 2 (SARS-CoV-2) Surface and Air Contamination in an Acute Healthcare Setting During the Peak of the Coronavirus Disease 2019 (COVID-19) Pandemic in London. Clin Infect Dis 73, e1870–e1877 (2021).

35. V. M. Corman et al., Detection of 2019 novel coronavirus (2019-nCoV) by real-time RT-PCR. Euro Surveill 25, 2000045 (2020).

36. A. Singanayagam et al., Characterising viable virus from air exhaled by H1N1 influenza-infected ferrets reveals the importance of haemagglutinin stability for airborne infectivity. PLoS Pathog 16, e1008362 (2020).

37. J. Zhou et al., Mutations that adapt SARS-CoV-2 to mink or ferret do not increase fitness in the human airway. Cell Rep 38, 110344 (2022).

38. L. Dupont et al., Neutralizing antibody activity in convalescent sera from infection in humans with SARS-CoV-2 and variants of concern. Nat Microbiol 6, 1433–1442 (2021).

39. J. Seow et al., Longitudinal observation and decline of neutralizing antibody responses in the three months following SARS-CoV-2 infection in humans. Nat Microbiol 5, 1598–1607 (2020).

40. K. Grehan, F. Ferrara, N. Temperton, An optimised method for the production of MERS-CoV spike expressing viral pseudotypes. MethodsX 2, 379–384 (2015).

41. C. P. Thompson et al., Detection of neutralising antibodies to SARS-CoV-2 to determine population exposure in Scottish blood donors between March and May 2020. Euro Surveill 25, (2020).

42. R. Hale et al., Development of a Multiplex Tandem PCR (MT-PCR) Assay for the Detection of Emerging SARS-CoV-2 Variants. Viruses 13, (2021).

